# Deep proteomics identifies shared molecular pathway alterations in synapses of schizophrenia and bipolar disorder patients and mouse model

**DOI:** 10.1101/2022.09.21.508852

**Authors:** Sameer Aryal, Kevin Bonanno, Bryan Song, D.R. Mani, Hasmik Keshishian, Steven A. Carr, Morgan Sheng, Borislav Dejanovic

## Abstract

Synaptic dysfunction is implicated in the pathophysiology of schizophrenia (SCZ) and bipolar disorder (BP). We used quantitative mass-spectrometry to carry out deep and unbiased profiling of the proteome of synapses purified from the dorsolateral prefrontal cortex of 35 cases of SCZ, 35 cases of BP, and 35 controls. Compared to controls, SCZ and BP synapses showed substantial and similar proteomic alterations. Network and gene set enrichment analyses revealed upregulation of proteins associated with autophagy and certain vesicle transport pathways, and downregulation of proteins related to synaptic, mitochondrial, and ribosomal function in the synapses of individuals with SCZ or BP. Some of the same pathways (e.g., upregulation of vesicle transport, downregulation of mitochondrial and ribosomal proteins) were similarly dysregulated in the synaptic proteome of mutant mice deficient in *Akap11*, a recently discovered shared risk gene for SCZ and BP. Our work provides novel biological insights into molecular dysfunction at the synapse in SCZ and BP and serves as a resource for understanding the pathophysiology of these debilitating neuropsychiatric disorders.

## Introduction

Schizophrenia (SCZ) and bipolar disorder (BP) are debilitating psychiatric disorders with respective lifetime prevalences of approximately 0.4-0.8% and 1-2% (McGrath et al., 2008; Palmer et al., 2022). BP is characterized by biphasic mood episodes of mania and depression, and SCZ by psychosis, negative symptoms, and cognitive impairment (Kahn et al., 2015; Vieta et al., 2018). Both diseases have substantial genetic components, with heritabilities estimated at over 60%, and many common variants of small effect making the largest overall contribution towards genetic risk (Ruderfer et al., 2018).

Despite their diagnostic classification as distinct disorders—and clear differences in typical presentation—SCZ and BP share some clinical and neuropathological features. Similar structural abnormalities and brain rhythm alterations have been observed in the two disorders (Hibar et al., 2016; Narayanan et al., 2014). Individuals with SCZ often exhibit mood disorders, and more than 50% of individuals with BP experience psychotic episodes in their lifetimes (Dunayevich and Keck, 2000; Hartman et al., 2019). The diagnosis of an individual patient can change over time from BP to SCZ or vice versa. These features suggest that the two diseases might lie on different parts of a psychosis spectrum.

More importantly, there is increasing evidence that SCZ and BP share common genetic risk factors. The two disorders display overlapping genetic susceptibility from common variation (genetic correlation = 60-70%) (Lee et al., 2019), as well as substantial cross-disorder relative risks (for parent/offspring: BP/BP: 6.4, BP/SCZ: 2.4; SCZ/BP: 5.2, SCZ/ SCZ: 9.9) (Lichtenstein et al., 2009). The polygenic risk score (PRS) for SCZ is associated with psychosis in BP, while the PRS for BP is associated with manic behavior in SCZ (Ruderfer et al., 2018). Of particular interest, recent exome sequencing in BP has identified *AKAP11* (*A-kinase anchoring protein 11*) as a large-effect risk gene shared with SCZ (combined odds ratio (OR) = 7.06, *P* = 2.83 × 10E−9)(Palmer et al., 2022; Singh et al., 2022). Human genetic and clinical evidence therefore points towards a significant overlap in the etiology and pathophysiology of SCZ and BP.

Studies of common genetic variants associated with SCZ have consistently implicated biological pathways related to synaptic function in disease risk (Ripke et al., 2014; Trubetskoy et al., 2022). Rare variant studies have also highlighted that synaptic dysfunction may play a causal role in at least a subset of SCZ (Fromer et al., 2014; Gulsuner et al., 2020; Singh et al., 2022). For instance, the largest exome-sequencing study of SCZ to date revealed significant enrichment of ultra-rare loss-of-function variants in genes related to synaptic organization and function, including *GRIN2A, GRIA3, and TRIO*, all of which encode well known synaptic proteins (Singh et al., 2022). Recent GWAS studies have shown that BP risk variants are also enriched in genes associated with synaptic function (Mullins et al., 2021). Transcriptomic analyses from post- mortem brains have also implicated synaptic pathways in pathophysiology of SCZ and BP (Jaffe et al., 2018; Zandi et al., 2022).

If synaptic dysfunction plays a key role in the pathophysiology of SCZ and BP—as suggested by human genetics—it is likely that this would be reflected in molecular changes in synapses of patients with these disorders. Synapses can be purified using standard biochemical fractionation (Bayés et al., 2011; Sheng and Hoogenraad, 2007), and unbiased proteomic examination of such synaptic fractions has been powerful for discovering pathogenic mechanisms of neurodegenerative disease (Dejanovic et al., 2018).

Here we report results of a large-scale, deep, and unbiased proteomic profiling of synaptic fractions purified from the dorsolateral prefrontal cortex (DLPFC) of individuals with SCZ and BP. For comparison, we also profiled the synaptic proteome of mice that harbor heterozygous and homozygous deletion in *Akap11*. Remarkably, we found that human SCZ and BP synapses exhibited highly similar changes in a number of proteins and molecular pathways, including pathways related to synaptic function, vesicle transport, mitochondrial respiration, and mRNA translation. Some of the shared pathway changes in SCZ and BP synapses—notably, ribosomal, mitochondrial respiration, and certain vesicle trafficking pathways—were also altered in the same direction in the synaptic proteome of mutant mice deficient in Akap11.

## Results

### Deep proteomic profiling of synaptic fractions of individuals with SCZ and BP

To examine how the protein composition of synapses is altered in SCZ and BP, we purified synaptic fractions from post-mortem dorsolateral prefrontal cortex (DLPFC; Brodmann’s Area 46) of a cohort of 35 individuals with SCZ, 35 with BP, and 35 matched controls (CTRL) and carried out MS-based quantitative proteomics of each individual sample using multiplexed tandem mass tag (TMT16) labeling (Fig. 1A).

**Figure 1:**
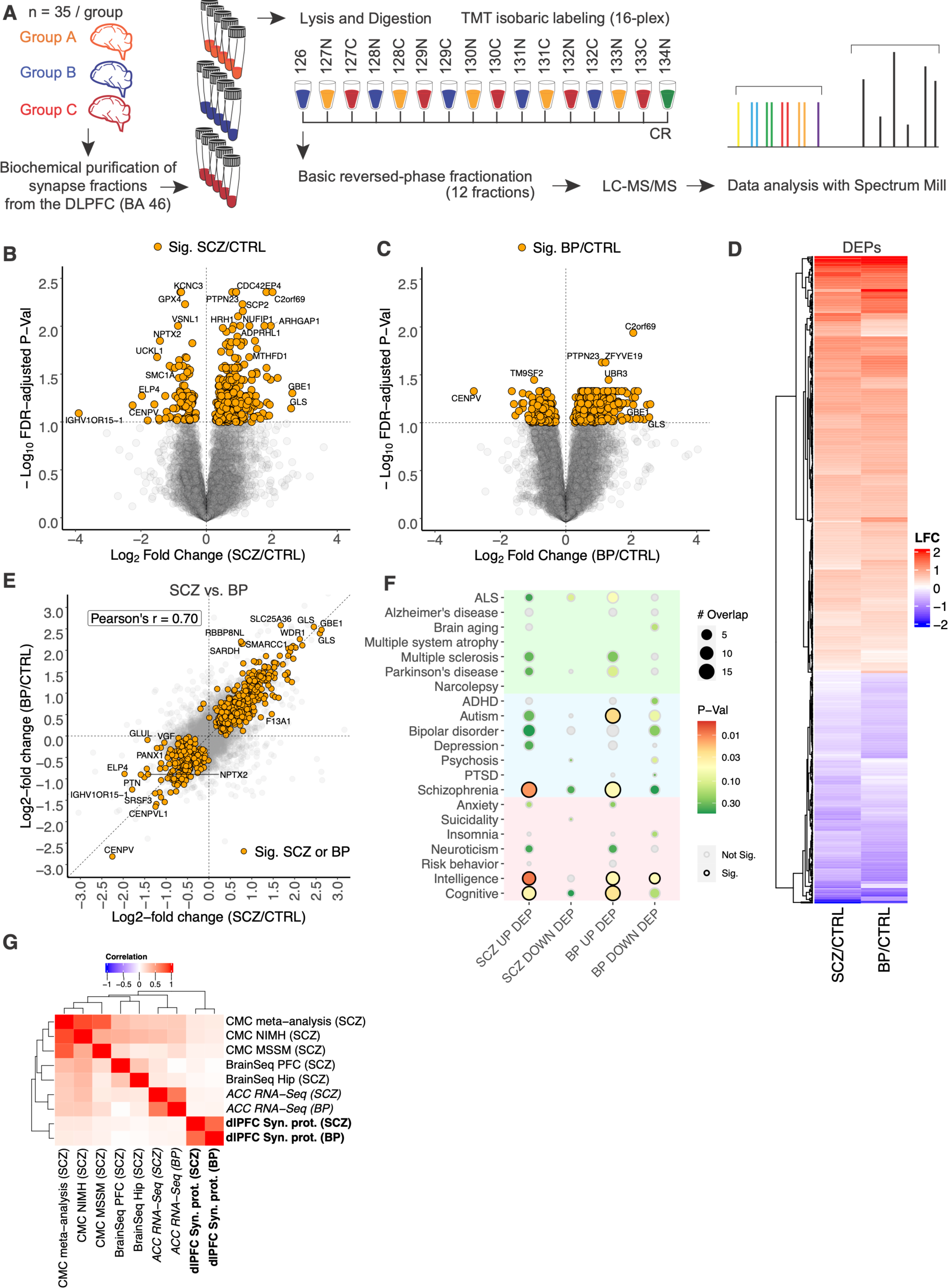
SCZ and BP synapses show substantial and similar proteomic changes. **(A)** Workflow used for sample processing and analysis. Synaptic fractions purified from the post- mortem DLPFC of groups A, B, and C (n=35 per group) were lysed and digested (see Methods). Five samples per group were included in each TMT16 plex along with a pooled common reference (CR). After labeling, samples were mixed and fractionated by basic reversed phase chromatography and fractions were analyzed by LC-MS/MS. Data was searched and summarized using Spectrum Mill MS Proteomics Software. Experiment and analysis were carried out blinded to group identity; samples from groups A, B, C were later revealed to correspond to BP, CTRL, and SCZ respectively. Examination of log2 fold-changes (LFCs) against FDR-adjusted p-values in **(B)** SCZ, and **(C)** BP synapses, compared to controls. DE proteins (DEPs; FDR- adjusted p-value < 0.1) are highlighted. **(D)** Heatmap showing LFCs in DEPs in either SCZ or BP synapses. **(E)** Evaluation of LFCs in SCZ synapses against those in BP. DEPs in either SCZ/CTRL or BP/CTRL are highlighted. **(F)** Bubble plot showing the number of DEPs that overlap with genes linked genetically to brain disorders or traits. The size of the bubble scales with the overlap; the color signifies the hypergeometric test nominal p-value, with those < 0.05 outlined to indicate significance. **(G)** Heatmap showing Pearson correlation coefficients evaluating LFCs in SCZ and BP synaptic proteomes against those in RNA-Seq from the anterior cingulate cortex (ACC) of the same cohort of individuals. Also shows Pearson’s r of SCZ and BP synaptic proteomics against those observed in large-scale RNA-Seq studies of SCZ, including those of the hippocampus (BrainSeq HIP) and PFC by the BrainSeq consortium, and the meta-, as well individual cohort analyses (MSSM and NIMH) of the PFC by the Commonmind consortium (CMC). Rows and columns are hierarchically clustered by Euclidean distance.

The average total protein yields of the synaptic fractions were not significantly different between groups, suggesting no gross changes in synapse density between CTRL, SCZ, and BP brain samples. (Fig. S1A). Of the 105 samples, 1 SCZ sample and 1 BP sample were excluded from further analysis due to MS-based evidence for blood contamination (Fig. S1I) and guidance from the brain bank, respectively (see Methods).

A total of 711,402 unique peptides that mapped to 8,996 proteins were identified, with each protein mapped by at least two unique peptides and detected in at least 20% of the samples (Supplementary Table T2; see Methods). We used a linear model that adjusted for covariates in the data to compute log2 fold-changes (LFCs) and carry out statistical testing for each identified protein (see Methods).

### Substantial and similar proteomic changes in SCZ and BP synapses

The synaptic proteomes of SCZ and BP samples were substantially different from those of matched controls (Fig. 1B, 1C). Using an FDR-adjusted P-value of < 0.1 as the threshold for differential expression (DE), we observed significant changes in the levels of 381 proteins (287 up, 94 down) in SCZ synapses (SCZ/CTRL; Fig. 1B), and 520 proteins (354 up, 166 down) in BP synapses (BP/CTRL; Fig. 1C).

Interestingly, 213 differentially expressed proteins (DEPs) were shared between SCZ and BP, with all DEPs altered in the same direction in SCZ and BP synapses (194 up, 19 down). In light of this overlap, we compared the LFCs of DEPs in either SCZ or BP synapses and observed highly correlated changes between the two disorders (Fig. 1D; Pearson’s r = 0.94). Notably, evaluation of the proteome-wide correlation in LFCs (i.e. all proteins) between SCZ and BP synapses also revealed a strong (Pearson’s r = 0.70) and significant correlation (P < 2.2e-16; Fig. 1E).

To exclude the possibility that using the same control group for both comparisons inflated the correlation observed between SCZ and BP synapses, we randomly split the CTRL samples in half and computed linear model LFCs for SCZ with half of the CTRL samples, and those for BP with the other half of CTRL samples. This process was repeated 10,000 times, and distributions of Pearson’s correlation coefficients and of p-values testing for positive correlations between the LFCs observed in SCZ/CTRL and BP/CTRL were examined. We observed a modal correlation coefficient of approximately 0.4 (Fig. S1B) and significant positive correlations between SCZ and BP LFCs in 98.7 % of the tests (implying a p-value of 0.01) (Fig. S1C). Thus, our analyses reveal that the synaptic proteomes in SCZ and BP feature unexpectedly similar alterations, suggesting a synaptic molecular pathology that is at least partly shared between the two disorders.

### Known and potential links between synapse DEPs and SCZ or BP

Neuronal pentraxin 2 (NPTX2) was reduced in both SCZ synapses (LFC = -1.42, FDR-adj. P = 0.0142) and to a lesser degree BP synapses (LFC = -0.896, FDR-adj. P = 0.109) (Fig. 1E; Fig. S1D). NPTX2, encoded by an immediate-early gene, is secreted by excitatory neurons and is important for maintaining excitatory homeostasis by regulating excitatory synapses on inhibitory interneurons (Xiao et al., 2021). Reduction of NPTX2 protein has been observed in cerebrospinal fluid (CSF) in SCZ as well as frontotemporal dementia (FTD) and Alzheimer’s disease (AD); CSF NPTX2 has therefore been proposed as a candidate biomarker for these diseases (Xiao et al., 2017; van der Ende et al., 2020; Xiao et al., 2021).

We also noted pronounced upregulation of C2orf69, a regulator of mitochondrial function (Lausberg et al., 2021; Wong et al., 2021), in both SCZ (LFC = 2.02, FDR-adj. P: = 0.004) and BP synapses (LFC = 2.06; FDR-adj. P = 0.012; Fig. 1B, 1C; Fig. S1E). Genome-wide and transcriptome-wide association studies (Gusev et al., 2018; Lee et al., 2019), as well as a genome-wide methylation quantitative trait loci study (Lin et al., 2018), have linked the *C2orf69* locus with increased risk for SCZ.

We next asked whether genes encoding DEPs in SCZ or BP synapses are enriched for GWAS ‘hits’ associated with SCZ, BP, and other neurological disorders or psychiatric traits. Hypergeometric tests revealed that DEPs upregulated in SCZ and BP synapses were nominally (unadjusted P < 0.05) enriched in genes linked genetically to SCZ risk (P = 0.01 (SCZ), 0.04 (BP); Fig. 1F; Fig. S1F). We also observed nominally significant enrichment of DEPs in SCZ and BP synapses in genes linked to intelligence and cognition. DEPs elevated in BP synapses were also over-represented in genes linked to autism. These genetic enrichment analyses add weight to the idea that DEPs might be causally involved in SCZ and BP, and that they play a role in cognitive function in humans.

### Synaptic proteome changes are not captured by transcriptome profiling

We wondered if the observed changes in level of specific proteins in the SCZ and BP synapses might be explained by changes in mRNA expression of these genes in the brain. To this end, we analyzed previously generated bulk RNA-Seq data (see Methods) from total homogenate of the anterior cingulate cortex (ACC) from the same cohort of individuals and compared the RNA-Seq versus synapse proteome LFCs. The correlations between RNA and synaptic protein alterations were weak in both SCZ and BP (Pearson’s r < 0.1; Fig. 1G; Fig. S1G, S1H). We also examined the synaptic proteome changes in SCZ and BP against previously published large-scale RNA-Seq studies of SCZ post-mortem brains and observed similarly weak correlations (Fig. 1G) (Collado-Torres et al., 2019; Hoffman et al., 2022). Notably, the RNA expression changes also did not replicate well across cohorts (Fig. 1G). Our analyses suggest that changes in synaptic protein levels are unlikely to be due to altered mRNA expression.

### Protein co-expression analysis identifies prominent reduction of canonical synaptic proteins and pathways in SCZ and BP

To gain a molecular systems-level understanding of changes in the synaptic proteome in SCZ and BP, we used weighted gene co-expression analysis (WGCNA), a computational method that categorizes genes and/or proteins with highly correlated expression in cases and controls into co-expression modules (Langfelder and Horvath, 2008). Over 8000 proteins were used to build the WGCNA network. We resolved 24 modules, or groups of proteins with highly similar expression patterns, from our synaptic protein co-expression network (Fig. 2). Enrichment analysis of proteins in the modules (see Methods) was used to identify the principal pathway(s) represented by each module. The number of proteins in the modules ranged from 39 to 1151; 973 proteins showed uncorrelated expression and were therefore placed in the M0 module which comprises proteins not assigned to any other modules.

**Figure 2:**
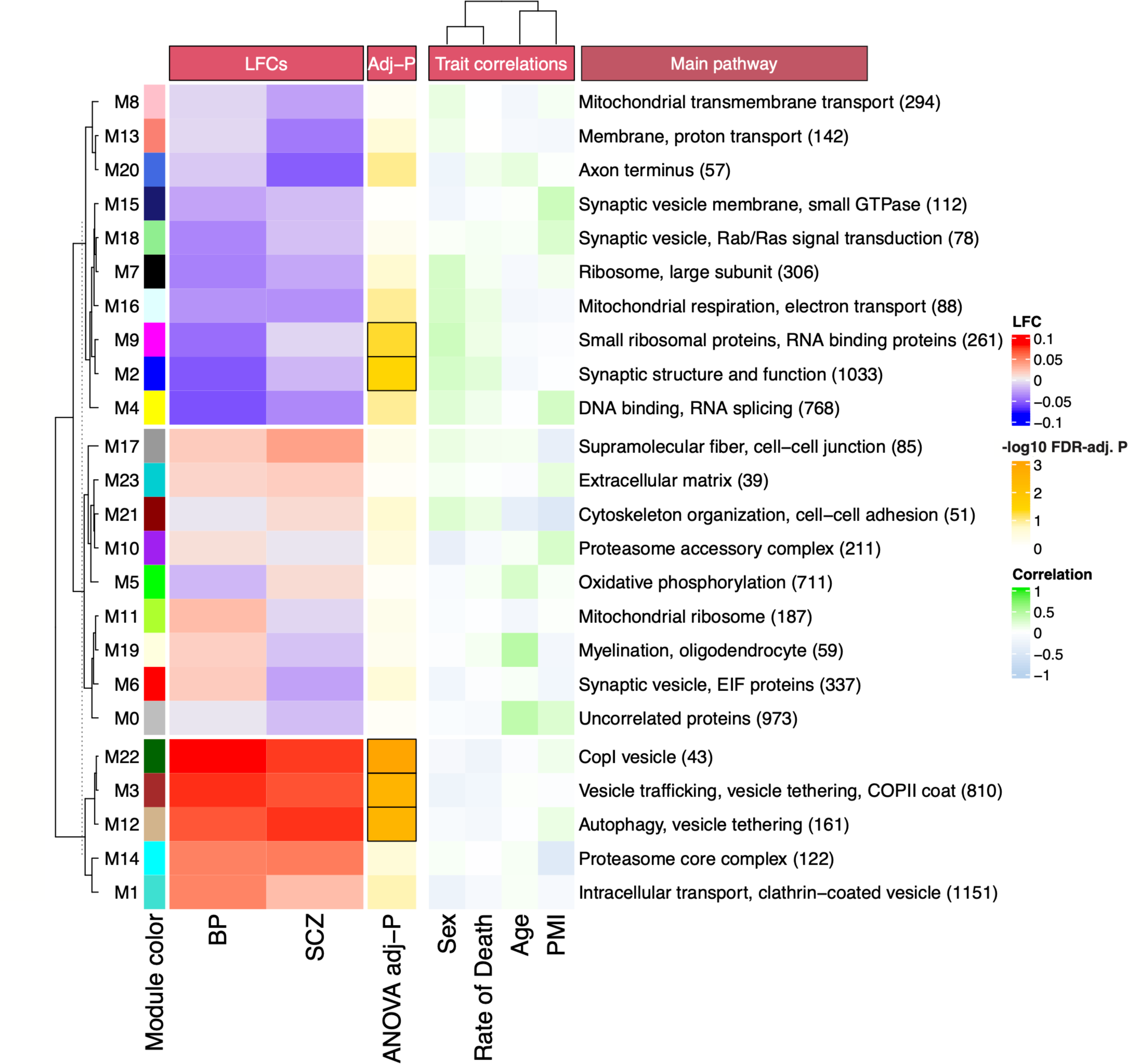
Synaptic protein co-expression network in SCZ and BP. Heatmap depicting the modules identified and their correlations with covariate traits in the synaptic protein co-expression network built using WGCNA. Each row represents a module; LFCs depict the change in module eigenprotein levels in SCZ and BP. Rows are hierarchically clustered by module eigenprotein LFCs in SCZ and BP, and the heatmap split into three based on k-means clustering of the same. The standard WGCNA module colors are shown. Also shown are the FDR-adjusted ANOVA p- values (-log10 transformed) testing for statistical difference in module eigenprotein levels across disease status; cells for modules with FDR-adjusted P < 0.1 are outlined. The primary biological pathway enriched in that module is also listed; in brackets are the number of the number of proteins in the module.

To examine which modules were associated with disease, we constructed a linear model of module eigenprotein (the first principal component of module protein expression) as a function of disease status, with covariates (sex, rate of death, age, and post-mortem interval (PMI)) included in the model to regress out their effects (Fig. 2). Examining the one-way analysis of variance (ANOVA) p-values of this model for disease status revealed that five modules were significantly changed in SCZ/BP (FDR-adjusted P < 0.1) (Fig. 2).

We observed significant reduction in module 2 (M2; FDR-adj. P = 0.041), which consisted of 1033 proteins, in SCZ and BP (Fig. 2, 3A). GO analysis of M2 revealed that the module was enriched in cellular components primarily related to the PSD organization and function, including ‘postsynaptic specialization, ‘glutamatergic synapse,’ and ‘postsynaptic membrane’ (FDR-adj. P < 1E-18 for all sets; Fig. 3B). Consistently, additional enrichment analysis with SynGO (Koopmans et al., 2019) confirmed the over-representation of protein sets related to post-synaptic organization, function, and signaling in M2, and further revealed the enrichment of presynaptic protein sets (Fig. S3A).

**Figure 3:**
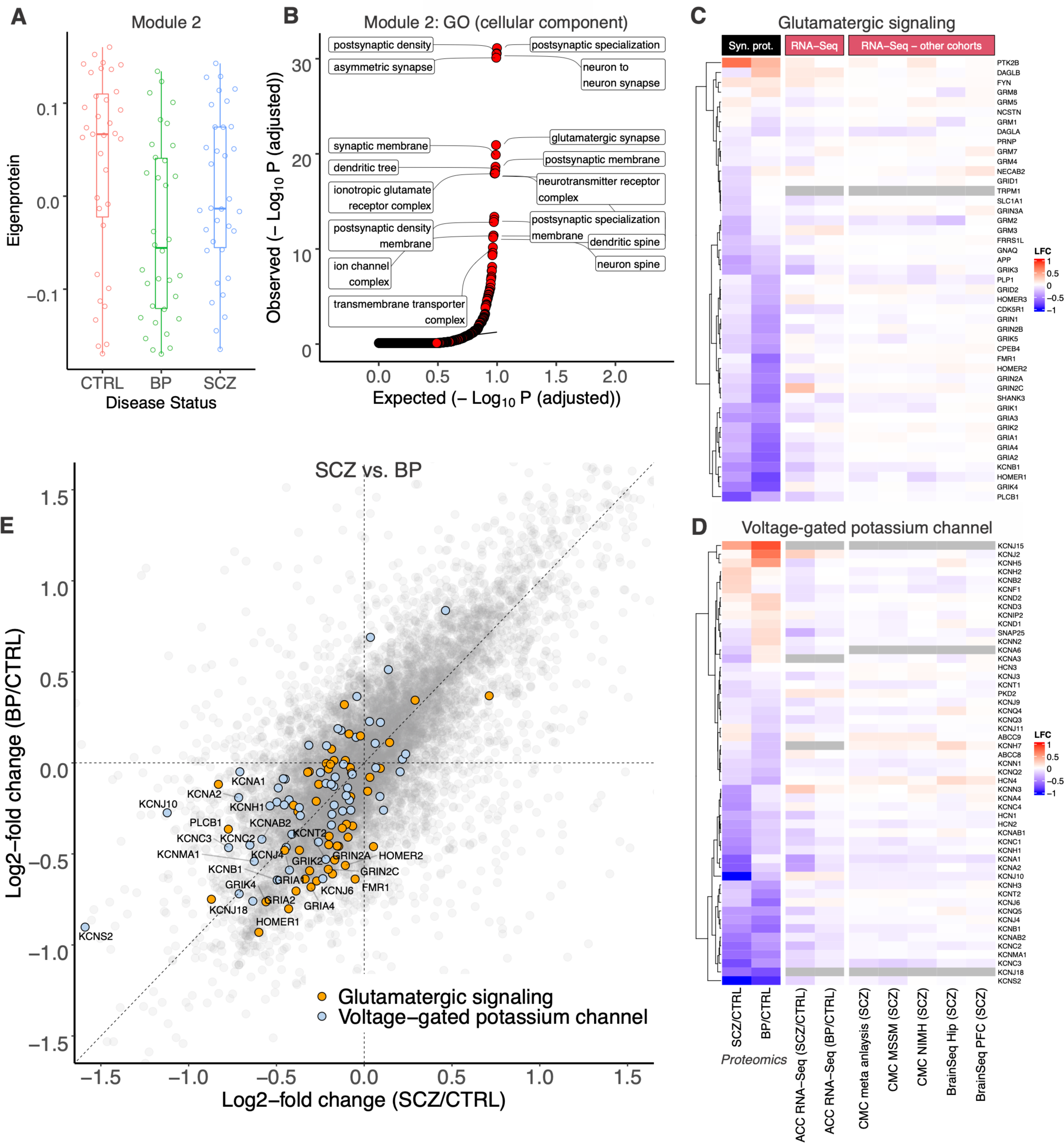
Core synaptic proteins are reduced in synapses from humans with SCZ and BP. **(A)** Module eigenprotein levels by disease status for module 2. **(B)** Q-Q plot examining cellular component GO-term enrichments in module 2. Heatmap of LFCs in all detected proteins in **(C)** the ‘glutamate receptor signaling pathway,’ and **(D)** ‘voltage-gated potassium channel’ gene sets in mSigDb. Each row represents an individual protein in the set. Also shown are the LFCs observed across multiple RNA-Seq studies. Rows are hierarchically clustered by synaptic proteome LFCs. **(E)** Evaluation of LFCs in SCZ against those in BP, with members of the ‘glutamate receptor signaling pathway,’ and ‘voltage-gated potassium channel’ highlighted.

Gene set enrichment analysis (GSEA) of the synaptic proteome-wide changes in SCZ/CTRL and BP/CTRL (see Methods) also corroborated these observations: Multiple protein sets related to synapses and synaptic function were reduced in BP and SCZ synapses, including ‘ion transmembrane transporter activity’ (FDR-adj. P = 1.05E-18 (SCZ), 1.46E-10 (BP)), ‘gated channel activity’ (FDR-adj. P = 1.38E-08 (SCZ), 4.63E-09 (BP)), and ‘dendrite membrane’ (FDR- adj. P = 9.59E-05 (SCZ), 3.60E-05 (BP)) (Supplementary Table T3). Notably, the ‘glutamate receptor signaling pathway’ protein set was significantly reduced in both SCZ and BP synapses, although only a few individual proteins of the set reached FDR-adj. p value < 0.1 (Fig. 3C, 3E; Fig. S3B, S3C), with BP synapses showing a larger reduction (normalized enrichment score (NES) =-2.4; FDR-adj. P=4.87E-06) than SCZ (NES=-1.68; FDR-adj. P=0.067). For instance, among the glutamatergic signaling set, there were significant reduction in Kainate receptor subunits GRIK2 and GRIK4 in BP synapses (Fig. S3C). SCZ synapses showed significant decreases in amyloid-precursor protein (APP) and the synaptic intracellular signaling molecule PLCB1 (Fig. S3B).

Interestingly, GSEA also revealed a striking reduction of proteins related to voltage-gated potassium channels (VGKC), with the magnitude of the reduction larger in SCZ (NES = -2.81; FDR-adj. P = 2.3E-11; Fig. 3D, 3E) than in BP synapses (NES = -2.03; FDR-adj. P = 0.02; Fig. 3D, 3E). Multiple VGKC proteins were significantly reduced in SCZ (n = 13; Fig. S3D) and BP synapses (n=6; Fig. S3E). Not only did SCZ and BP synapses show changes in the same pathways (glutamatergic signaling and VGKCs, Fig. 3C, 3D) but they displayed correlated changes in the individual proteins of those pathways (Fig. 3E).

Examining intramodular hub-proteins, which are the proteins most correlated with the module eigenprotein, revealed the Regulating synaptic membrane exocytosis protein 1 (RIMS1; LFC = -0.35 (SCZ), -0.63 (BP)) as the protein ‘anchor’ of M2. Interestingly, GWAS loci at *RIMS1*, which regulates synaptic vesicle exocytosis, have been associated with increased risk for SCZ (Lee et al., 2019; Ripke et al., 2014) and BP (Stahl et al., 2019). Overall, our data show that key pre- and postsynaptic proteins and cation channels, especially glutamate receptors and VGKC, are reduced in both SCZ and/or BP synapses.

### Vesicular transport and intracellular trafficking related pathways are elevated in SCZ and BP synapses

The most significantly upregulated WGCNA module in BP/SCZ synapses was module 22 (M22), which consists of 43 proteins (Fig. 4A), almost all of which were increased in SCZ and BP synaptic proteomes (Fig. S4A). GO analysis of cellular components indicated that proteins from the coat protein complex I (COPI) and related GO terms were particularly over-represented in M22 (Fig. 4B). COPI protein-coated vesicles are best characterized as mediating retrograde transport from the trans-Golgi network to the cis-Golgi network and endoplasmic reticulum (ER) (Popoff et al., 2011). M22 was anchored by the COPI protein Coat complex subunit beta-1 (COPB1) and includes multiple other COPI vesicle coat proteins, such as COPI Coat complex subunits alpha (COPA), beta-2 (COPB2), gamma-1 (COPG1), gamma-2 (COPG2), and Archain 1 (ARCN1) (Fig. S4A).

**Figure 4:**
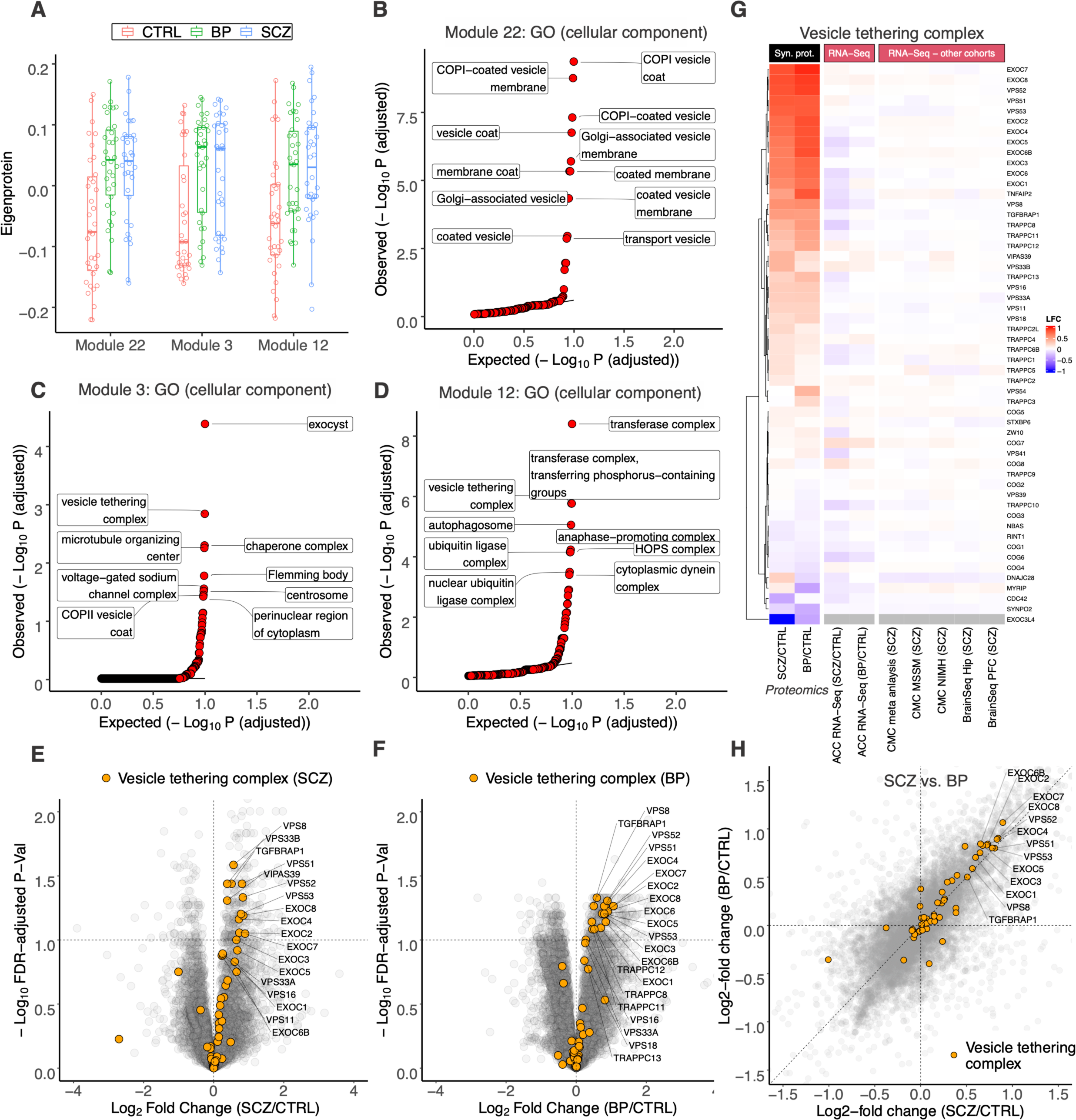
Proteins related to vesicle tethering and trafficking are upregulated in SCZ and BP synapses. **(A)** Boxplots showing module eigenprotein levels stratified by disease status for modules 22, 3, and 12. Q-Q plot examining cellular component GO-term enrichments in **(B)** module 22, **(C)** module 3, and **(D)** module 12. Volcano plot displaying LFCs against FDR-adjusted p-values in **(E)** SCZ synapses, and **(F)** BP synapses; members of the ‘vesicle tethering complex’ set in mSigDb are highlighted. **(G)** Heatmap showing LFCs in vesicle tethering complex proteins in SCZ and BP synapses; alterations (LFCs) in RNA expression of the same proteins observed across RNA-Seq studies are also shown. Heatmap is hierarchically clustered by SCZ and BP synaptic proteome LFCs. **(H)** Evaluation of LFCs in SCZ synapses against those in BP, with proteins in the ‘vesicle tethering complex’ set highlighted.

Interestingly, we found that module 3 (M3), the second most significantly upregulated module (Fig. 4A), was enriched in COPII vesicle coat proteins, a distinct set of proteins from those that coat COPI vesicles (Fig. 4C). COPII-coated vesicles mediate the anterograde transport of newly synthesized proteins from the ER to the Golgi apparatus (Miller and Schekman, 2013). The COPII proteins SEC31A, SEC23A, SEC13, and SEC24C were significantly increased in SCZ and/or BP synapses (Fig. S4B, S4C). These findings indicate that proteins involved in both anterograde and retrograde ER-to-Golgi vesicle trafficking are significantly and similarly altered in SCZ and BP synapses. The most significantly enriched cellular component in M3, however, was the ‘exocyst complex’, a subset of the larger – also highly enriched – ‘vesicle tethering complex’ superset (Fig. 4C). Vesicle tethers are protein complexes that mediate the initial attachment of a transport vesicle to its target membrane prior to fusion (An et al., 2021). For instance, the exocyst complex facilitates the spatial targeting and tethering of post-Golgi vesicles to the plasma membrane. The Golgi-associated retrograde protein (GARP) complex, which is involved in the transport of endosomes from the plasma membrane to the Golgi (Fröhlich et al., 2015), was also enriched in M3 (Supplementary Table T4).

Interestingly, module 12 (M12), the final significantly upregulated module in SCZ/BP synapses (Fig. 4A), was also enriched in multiple subsets of the vesicle tethering superset (Fig. 4D; Supplementary Table T4). While M3 captured the elevations in exocyst and GARP complexes, M12 captured those in the TRAPP III (transport protein particle III), and the homologous CORVET (class C core vacuole/endosome tethering) and HOPS (homotypic fusion and protein sorting) complexes. The CORVET and HOPS complexes tether endosome-endosome and endosome-lysosome fusion respectively (Nickerson et al., 2009); and the TRAPP complex is a tethering factor for COPII vesicles (Tan et al., 2013).

Consistent with the findings from the WGCNA, GSEA of proteomic changes in SCZ and BP synapses indicated that ‘vesicle tethering complex’ was one of the most significantly increased protein sets in both SCZ and BP synapses (Supplementary Table T3). Examining the alterations in the proteins comprising this set revealed significant elevation of multiple exocyst- (e.g., EXOC2, -7, -8), GARP- (e.g., VPS51, -52, -53), CORVET/HOPS- (e.g., VPS8, -16, TGFBRAP1), and TRAPP III-complex proteins (e.g., TRAPPC8, -11, -12) in SCZ and/or BP synapses (Fig. 4E, 4F). The magnitude and direction of changes in vesicle tethering proteins were strikingly correlated across BP and SCZ synapses (Fig. 4G, 4H).

While protein sets related to vesicle tethering are some of the most enriched GO terms in M3 and M12, it should be noted that these modules, especially M3 (containing 810 proteins), consist of a large number of proteins, and consequently they likely capture networks related to multiple forms of vesicular trafficking and related biological pathways. In fact, the top three hub proteins of the M3 are the Phospholipase A-2-activating protein (PLAA), which is involved in trafficking of synaptic membrane proteins to late endosomes in a ubiquitin-dependent manner (Hall et al., 2017); Guanine nucleotide-binding protein-like 1 (GNL1), which may have a putative role in activating the ARF GTPases that mediate membrane trafficking (Teh and Moore, 2007); and Dynamin-1 (DNM1), a GTPase with well-known roles in membrane remodeling and vesicular fission/trafficking, including at synapses (Ferguson and De Camilli, 2012). We additionally noted that M3 eigenproteins were strongly negatively correlated with those from M2 (Pearson’s r = - 0.82), suggesting that at an individual sample level, regardless of disease status, a reduction in synaptic proteins is associated with an increase in certain vesicle trafficking proteins at the synapse (Fig. S4D).

Taken together, our findings suggest that multiple distinct mechanisms of vesicular transport and intracellular trafficking, including those involved in retrograde and anterograde Golgi-to-ER trafficking, post-Golgi vesicular tethering, trafficking, and endosomal transport are altered at SCZ and BP synapses. These protein changes at the synapse seem not to be driven by transcriptional changes (Fig. 4G, Fig. S4A).

### Autophagy-related pathways are increased in SCZ and BP synapses

Interestingly, GO analysis of biological processes enriched in M12 (which is upregulated in both SCZ and BP) revealed autophagy to be the most enriched protein set in the module (Fig. 5A), consistent with the strong over-representation of the ‘autophagosome’ cellular component in this module (Fig. 4D). Enrichments for ‘macroautophagy’ and ‘autophagosome assembly’ gene sets were also observed in the M3 module (Fig. S5A; Supplementary Table T4). It is known that COPII as well as TRAPPIII complex proteins, which are both elevated in SCZ/BP synapses (Fig. 4G), can promote autophagy during nutrient stress (Tan et al., 2013; Jeong et al., 2018; Harris et al., 2021). These findings suggest that autophagy may be activated at SCZ and BP synapses.

**Figure 5:**
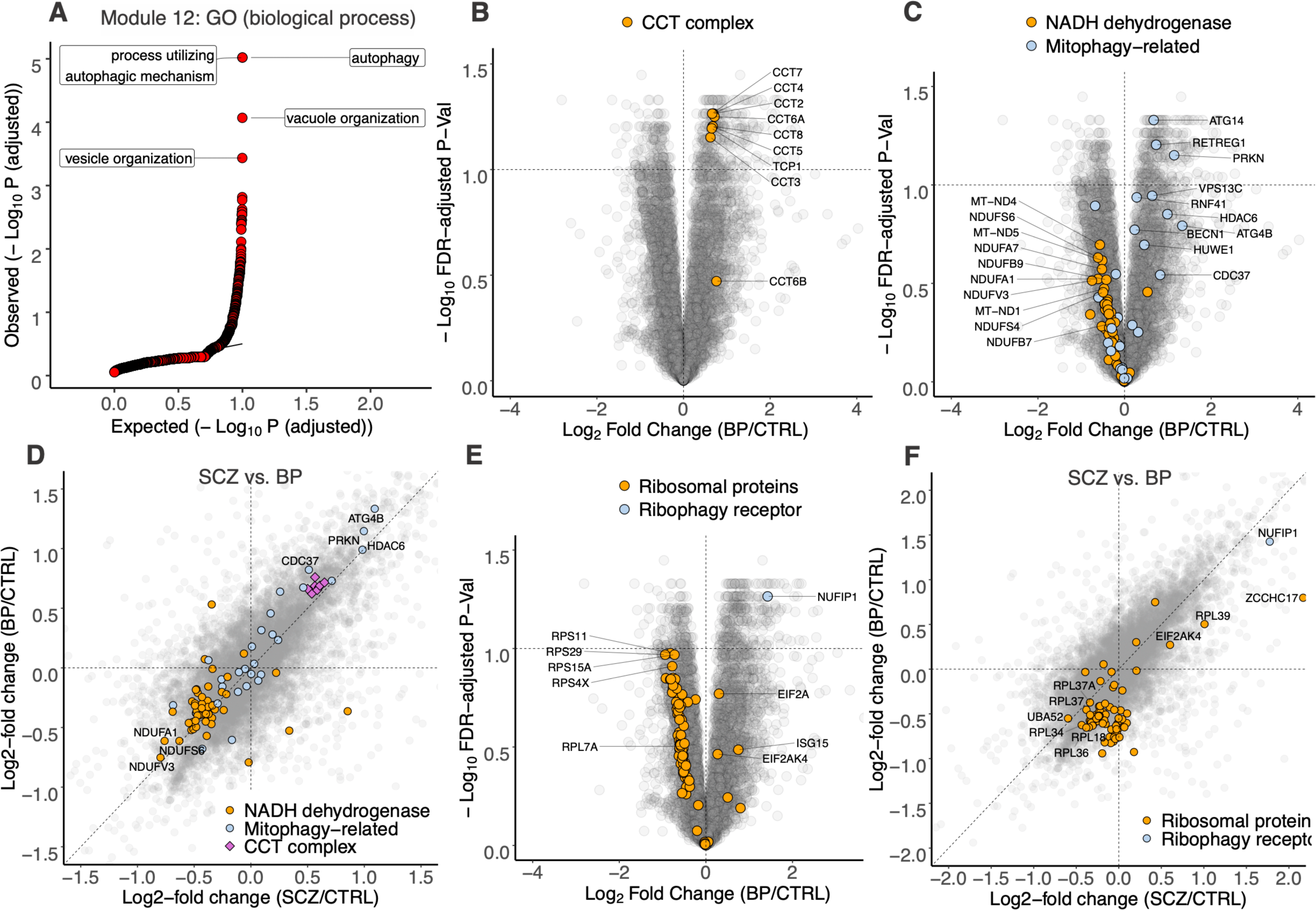
Autophagy pathways are elevated in SCZ and BP synapses. **(A)** Q-Q plot showing biological processes enriched in module 12. Volcano plot showing alterations in BP synapses highlighting all detected proteins in the **(B)** ‘chaperonin-containing T (CCT) complex,’ and **(C)** ‘NADH dehydrogenase complex,’ and ‘mitophagy’ gene sets in mSigDb. **(D)** Alterations in SCZ synapses compared to those against BP, with proteins related to mitochondrial respiration (‘NADH dehydrogenase complex’), ‘mitophagy’, and CCT complex gene sets highlighted. **(E)** Volcano plot highlighting the alterations in ribosomal proteins (‘cytosolic ribosome’ gene set), and the ribophagy receptor NUFIP1 in BP synapses. **(F)** Evaluation of LFCs in SCZ against those in BP synapses; ribosomal proteins, and the ribophagy receptor NUFIP1 are colored.

GO analysis of upregulated DEPs revealed (macro)autophagy as one of most significantly enriched pathways in BP and SCZ (Fig. S5B; S5C). It also revealed a striking increase in the Chaperonin-containing TCP-1 (CCT, also known as TRiC) complex in BP (Fig. 5B) and SCZ synapses (Fig. S5D), including CCT2, -4, -6, -8 etc. (Fig. 5B, 5D; Fig. S5D). Previous work has shown that reduction in CCT complex proteins impairs autophagy (Pavel et al., 2016) and a recent study has further identified CCT2 as a receptor for aggrephagy (Ma et al., 2022). The robust increase in the CCT complex therefore provides additional evidence suggesting an increase in autophagy at SCZ and BP synapses.

Multiple well-known proteins involved in mitophagy, such as Parkin (PRKN; an E3 ubiquitin ligase that promotes mitophagy) and ATG14 (which targets phosphatidylinositol 3-kinase complex I to the phagophore assembly site) were also notably upregulated in BP synapses (Fig. 5C), and more modestly so in SCZ synapses (Fig. S5E). Interestingly, this elevation in mitophagy-related proteins was accompanied by a reduction in protein sets related to outer- (e.g., ‘Outer Mitochondrial Membrane Protein Complex’; FDR-adj. P = 0.02 (SCZ), 0.007 (BP)), and more prominently, inner-mitochondrial membranes (e.g., ‘Inner mitochondrial membrane protein complex’, FDR-adj. P < 2E-8 SCZ and BP), particularly those mediating respiration and electron transport (e.g., ‘NADH dehydrogenase complex’, FDR-adj. P = 1.47E-6 (SCZ), 6.7E-5 (BP); Fig. 5C, 5D; Fig. S5E).

We also observed strong upregulation of NUFIP1, the ribosome receptor for starvation- induced ribophagy, and a more modest increase in the starvation-induced autophagy- and ER stress-activator GCN2 (EIF2AK4), in SCZ (Fig. S5F) and BP synapses (Fig. 5E, 5F) (Ravindran et al., 2014; Tallóczy et al., 2002; Wyant et al., 2018). These increases in NUFIP1 and EIF2AK4 were accompanied by a widespread reduction in ribosomal proteins (Fig. 5E; Fig. S5F; ‘Structural constituent of ribosome’, FDR-adj. P= 0.08 (SCZ), 3.6E-8 (BP)). Overall, our data suggests that autophagy processes (including ribophagy and mitophagy) may be induced at the synapse in SCZ and BP.

The respiration and electron transport chain-related proteins, small ribosomal proteins, and large ribosomal proteins were captured by modules 16, 9, and 7 respectively; while all three modules were reduced in BP and SCZ synapses, only module 9 (M9) was statistically significant (ANOVA FDR-adj. P = 0.12 (M16); 0.06 (M9); 0.23 (M7) respectively; Fig. 2). M9 was also enriched in RNA-binding proteins (RBPs) related to translation initiation; in fact, its hub protein GIGYF2 (GRB10 Interacting GYF Protein 2; LFC= -0.17 (SCZ), -0.34 (BP)) functions to repress the translation of defective mRNAs (Hickey et al., 2020). Intriguingly, GWAS loci at *GIGYF2* have been associated with increased risk for SCZ (Ripke et al., 2014; Lam et al., 2019; Trubetskoy et al., 2022).

*Comparison of synapse proteome alterations in human SCZ and BP with Akap11 mutant mice* Exome sequencing has identified *AKAP11* as a shared risk gene for BP and SCZ (Palmer et al., 2022; Singh et al., 2022). We recently showed that *Akap11* mutant mice have abnormal features in electroencephalogram (EEG) recordings (Herzog et al., 2022). Furthermore, AKAP11, along with other protein products of SCZ risk genes, was present in the human synapse proteome (Fig. S6A). Thus, we hypothesized that *Akap11* mutant mice might be a suitable (and understudied) SCZ/BP animal model to study the molecular pathomechanisms at the synapse. We therefore profiled the synapse proteome from cerebral cortex of 1-month-old male wild-type (WT), heterozygous (Het) and homozygous (KO) *Akap11* mice (n=5 per genotype) and compared the changes found in *Akap11* mutant mice with those observed in human SCZ and BP.

Proteome-wide, the LFCs of proteins in *Akap11* Het and KO synapses (vs. WT) were strongly and highly significantly correlated with each other, though perhaps with a greater magnitude of change in KO (Pearson’ r = 0.75, P < 2.2e-16; Fig. S6B). The LFCs of *Akap11* Het or KO synapses showed little correlation with those of SCZ or BP synapses, at least at the proteome-wide level (Fig. 6A). Strikingly, however, by GSEA the gene sets that were significantly altered in Akap11 Het and KO mouse synapses were similar to those affected in SCZ and BP human synapses (in direction and enrichment score) across all four comparisons (Fig. 6B). Vesicle transport and intracellular trafficking pathways, which were among the most strongly upregulated molecular pathways in SCZ and BP synapses, were also among the most enriched gene sets in *Akap11* mouse Het and KO synaptic proteomes (Fig. 6B, Supplementary Table T5). Similarly, pathways associated with ion transport, ribosomes, and mitochondrial respiration were strongly reduced in *Akap11* Het and KO mouse synapses, concordant with the changes observed in SCZ and BP human synapses (Fig. 6B). For example, comparing the changes in the vesicle tethering complex gene set and in the ‘NADH dehydrogenase’ gene set in *Akap11* Het and KO synapses against those in SCZ and BP synapses confirmed that despite the overall poor correlation at the whole-proteome level, these same pathways were altered in synapses from Akap11-deficient mice and SCZ/BP subjects (Fig. 6C; Fig. S6C, S6D, S6E). The enrichment of the vesicle tethering complex in Akap11-deficient synapses was driven by strong changes in a subset of tethering proteins—primarily the Conserved Oligomeric Golgi (COG) complex proteins (e.g., COG3, -6, -8), thought to mediate tethering of COPI vesicles—as opposed to the large alterations in exocyst- (e.g., EXOC7, -8), GARP- (e.g., VPS51, -52, -53), and CORVET/HOPS- complex proteins (e.g., VPS8, -16) that drive the enrichment of vesicle tethering complex in SCZ and BP synapses (Fig. 6C; Fig. S6C, S6D, S6E). Thus, while there are species-specific DEP alterations in the synapse proteome, Akap11-deficient mouse synapses recapitulate many of the major pathway changes observed in SCZ and BP synapse proteomes, supporting the notion that these common altered pathways (such as vesicle trafficking, vesicle tethering, mitochondrial respiration) might play a pathophysiologic role in SCZ and BP.

**Figure 6:**
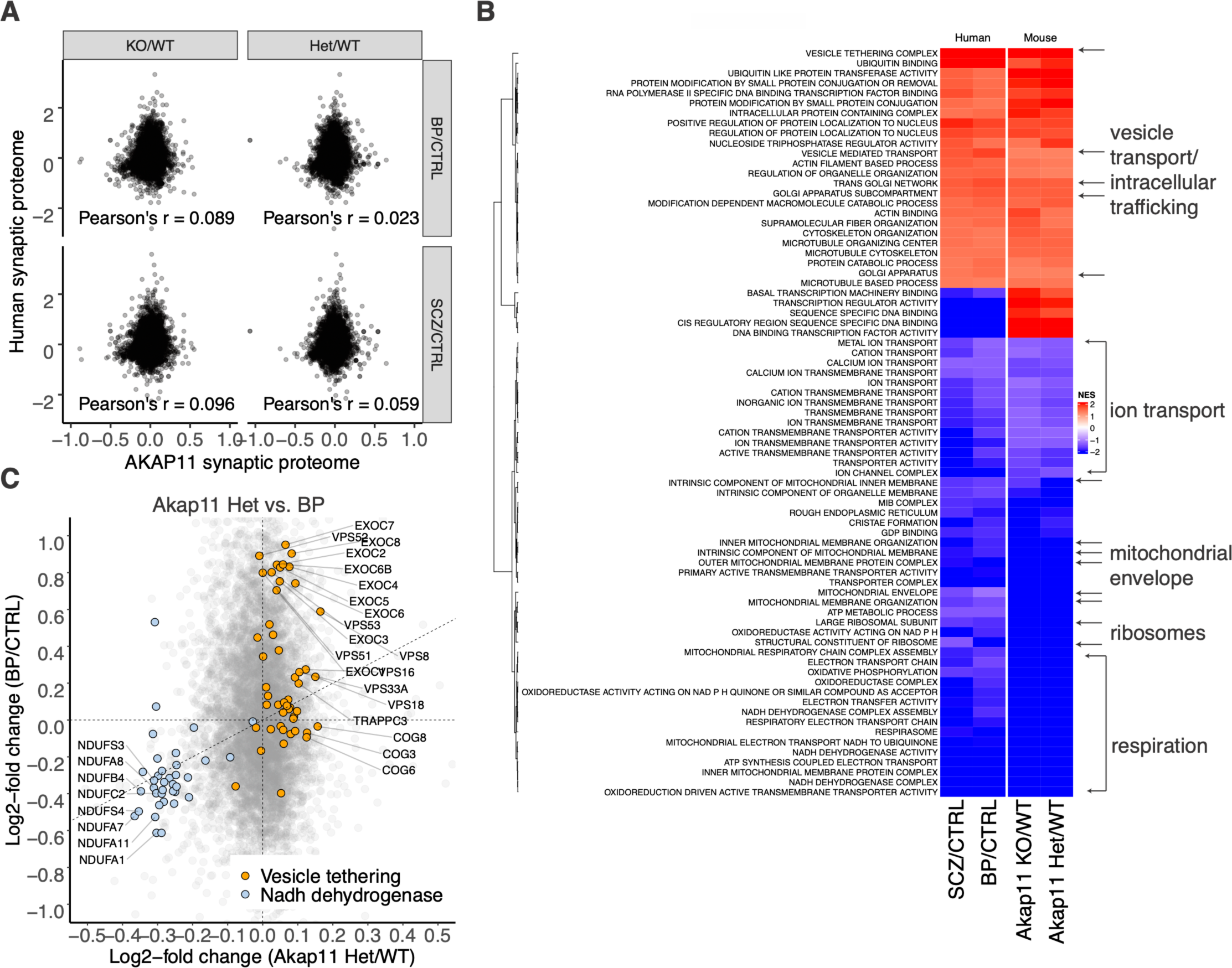
Comparison of alterations in SCZ and BP synapses with those in *Akap11* mice. **(A)** Evaluation of the cortical synaptic proteome LFCs in *Akap11* Het and KO mice against those observed in synapses purified from post-mortem DLPFC of individuals with SCZ and BP. **(B)** Heatmap showing gene sets that are significantly changed (FDR-adjusted p < 0.1) across all of SCZ, BP, *Akap11* Het, and *Akap11* KO synapses. **(C)** Examination of alterations in the synaptic proteome of *Akap11* Het mutants against those observed in post-mortem brains from individuals with BP; mSigDB gene sets related to mitochondrial respiration (‘NADH dehydrogenase complex’), and vesicle trafficking (‘vesicle tethering complex’) are colored.

## Discussion

Our study reveals pronounced molecular changes in synapses purified from postmortem DLPFC of humans with SCZ and BP and cerebral cortex of Akap11-deficient mice. In comparison to previously published synapse proteomes from either SCZ or BP patients (Föcking et al., 2015, 2016; MacDonald et al., 2020), our study (i) analyzes the SCZ and BP synapse proteome of the human DLPFC—a brain region of critical importance for both disorders (Sakurai et al., 2015; Abé et al., 2022); (ii) systematically compares SCZ and BP synapse proteomes against each other as well as against a genetic mouse model of both diseases; (iii) does so at unprecedented depth of protein- and protein network-identification, and (iv) features large numbers of samples comparable with or substantially larger than those in previous studies. Our unbiased deep proteomics datasets and systems-biology analyses therefore provide unprecedented resources and novel insights for understanding the proteins and protein networks associated with SCZ and BP.

One important and surprising finding is that SCZ and BP synapse proteomes exhibit similar differences compared to controls, suggesting that common molecular pathomechanisms act at the synapse for these two psychiatric disorders. Our proteomics data therefore lends biochemical support to the idea that SCZ and BP are related illnesses lying on a psychosis spectrum. In addition, we identified remarkably overlapping pathway changes in synapses from humans with SCZ/BP and mice deficient in *Akap11*, a shared risk gene for SCZ and BP. These findings highlight the utility of unbiased inter-species proteomic profiling of synapses—especially in human genetics-validated animal models—to identify protein (network) changes potentially involved in the pathophysiology of SCZ and BP.

Synapse dysfunction has long been considered as a mechanism underlying SCZ and BP pathophysiology, and is implicated by human genetics and gene expression studies (Jaffe et al., 2018; Mullins et al., 2021; Singh et al., 2022; Trubetskoy et al., 2022; Zandi et al., 2022). We observed reductions in a variety of canonical PSD proteins in SCZ and BP synapses, including glutamate receptors (AMPA, NMDA, and Kainate receptors), glutamatergic signaling pathway molecules (e.g., PLCB1, APP), and postsynaptic scaffolding proteins (e.g., HOMER1 and SHANK3), which supports the large body of evidence implicating glutamatergic receptor signaling impairment as a key pathomechanism in SCZ and BP. Interestingly, we previously showed that canonical PSD proteins are reduced in Tau-P301S mice (a mouse model of tauopathies like FTD and AD (Dejanovic et al., 2018)), indicating that there might be pathophysiological mechanisms between tauopathies and neuropsychiatric diseases that converge on core PSD proteins.

More unexpectedly, our analysis identified a significant increase in a number of protein trafficking and autophagy networks. We found evidence that (macro)autophagy processes, including mitophagy, ribophagy, and aggrephagy, may be elevated in SCZ and BP synapses. Autophagy is often stimulated under a variety of stress conditions such as nutrient/energy stress, ER stress, and redox stress (Murrow and Debnath, 2013). At the synapse, the autophagic machinery is induced by long-term depression leading to the degradation of core PSD proteins (Compans et al., 2021; Kallergi et al., 2022). The upregulated autophagy processes may therefore reflect cellular stress and contribute to changes of synapse composition in SCZ and BP. Increased mitophagy and ribophagy could, at least in part, explain the decrease in mitochondrial and ribosomal proteins in synapses of SCZ/BP patients and Akap11-deficient mice. Among other biological processes, synaptic mitochondria provide the energy supply for local translation during synaptic plasticity (Li et al., 2004; Rangaraju et al., 2019). Thus, reduced synaptic mitochondria content and decreased ribosomal protein levels suggest that synaptic protein biosynthesis (and synaptic plasticity) might be impaired in SCZ and BP, as well as in Akap11-deficient mice. Interestingly, previous SCZ and BP synapse proteome studies have also reported a reduction in mitochondrial proteins (Föcking et al., 2015, 2016; MacDonald et al., 2020), suggesting that bioenergetic abnormality at the synapse is a consistent and therefore relevant pathomechanism. There was very little correlation between the protein changes at the synapse and transcriptome changes in the cortex, implying that protein alterations at the synapse are due to changes in protein turnover (and/or protein subcellular distribution) rather than driven by transcriptional regulation. The putative increase in aggrephagy (as implied by upregulation of CCT complex proteins) could be associated with aberrant proteostasis, which might be triggered by cellular stress and lead to protein misfolding in SCZ/BP synapses. Consistent with this notion, increased protein insolubility and elevated ubiquitination—albeit at more subtle degrees than in neurodegenerative diseases—have been reported in SCZ and BP brains (Bradshaw and Korth, 2019; Leliveld et al., 2008; Nucifora et al., 2019).

The surge in the homologous CORVET and HOPS tethering complexes furthermore indicates that the endo-lysosomal pathway is reshaped in SCZ/BP synapses. While CORVET is generally required for fusion and maturation of early endosomes, HOPS mediates the fusion of late endosomes and autophagosomes with lysosomes (van der Beek et al., 2019). Furthermore, the highly elevated COPI and COPII mediate cargo transport between Golgi and ER, which is critical for autophagosome formation (Razi et al., 2009). The changes in intracellular trafficking pathways add support to the inference that (macro)autophagy-related pathways are increased in SCZ/BP synapses.

Furthermore, the prominent increase in vesicle tethering and forward and retrograde secretory trafficking proteins suggests that delivery and recycling of cargo is induced at the SCZ/BP synapse. For example, elevated TGFBRAP1/VPS3 and VPS8 indicate that transport to recycling endosomes might be induced at the SCZ/BP synapse (Jonker et al., 2018). The increase in exocyst complex and other vesicle tethering complexes might be particularly relevant for delivery and recycling of transmembrane proteins at synaptic sites, including AMPA, NMDA and potassium channel subunits (Broutman and Baudry, 2001; Ma and Jan, 2002; Sans et al., 2003). Despite the limited synaptic proteome-wide correlation in LFCs between Akap11 mutant mice and human SCZ/BP, we discovered that intracellular trafficking and vesicle tethering were also among the most highly induced pathways in synapses from Akap11-deficient mice. Further studies are needed to investigate how the alterations in protein and vesicle trafficking, autophagy, mitochondrial and protein translation pathways are contributing to disease pathophysiology.

Mining the DEPs in our dataset, especially those of secreted proteins, might yield promising fluid biomarkers for diagnosis and/or patient stratification of SCZ/BP. One of the most significantly decreased proteins in SCZ, and to a lesser degree in BP synapses, was NPTX2, an activity-induced secreted protein expressed by neurons (Chang et al., 2010). NPTX2 has been recently shown to be significantly reduced in the CSF of SCZ patients (Xiao et al., 2021). Nptx2 mediates homeostatic upscaling of AMPA receptors at excitatory synapses on parvalbumin interneurons and Nptx2-deficient mice exhibit behavioral deficits reminiscent of SCZ (Pelkey et al., 2015; Xiao et al., 2021), suggesting that NPTX2 might contribute to disease pathogenesis.

An obvious limitation of our study is the use of post-mortem human brain tissue. Although we attempted to control for confounding factors in our analysis, we cannot exclude that synapse integrity, and thereby the biochemical isolation, might have been impacted in some samples. Furthermore, we studied only patients with established (chronic) SCZ and BP, many of which took CNS medications that might affect synapse structure and function. While we cannot distinguish primary pathogenic changes from secondary compensatory or drug-induced changes in the synapse proteome of SCZ and BP disorders, we are encouraged that in our human-mouse comparative analysis, Akap11-deficient mice showed changes in several common pathways (such as membrane trafficking, vesicle tethering, mitochondrial respiration, protein synthesis) that were similarly altered in SCZ and BP synapses. The overlap in synapse proteomics changes enhances confidence that the human dataset is capturing, at least in part, the molecular features that are relevant to SCZ and BP disease mechanisms.

Our large-scale proteomics analysis provides an unbiased view of the composition of the SCZ and BP synapse proteome at unprecedented depth, and does so in comparison with a mouse model of SCZ and BP that is based on human genetics. The limited overlap between synaptic proteome and global transcriptome highlights the importance of analyzing proteins in addition to transcriptomics for understanding mechanisms of disease. Our data sheds new light on SCZ and BP pathophysiology, serves as a roadmap for future studies in human and mouse models, and prioritizes protein pathways and networks for biomarker discovery and disease-modifying therapeutic development.

## Supporting information

Supplemental Tabel T1

Supplemental Tabel T2

Supplemental Tabel T3

Supplemental Tabel T4

Supplemental Tabel T5

## Figure legends

**Figure S1:**
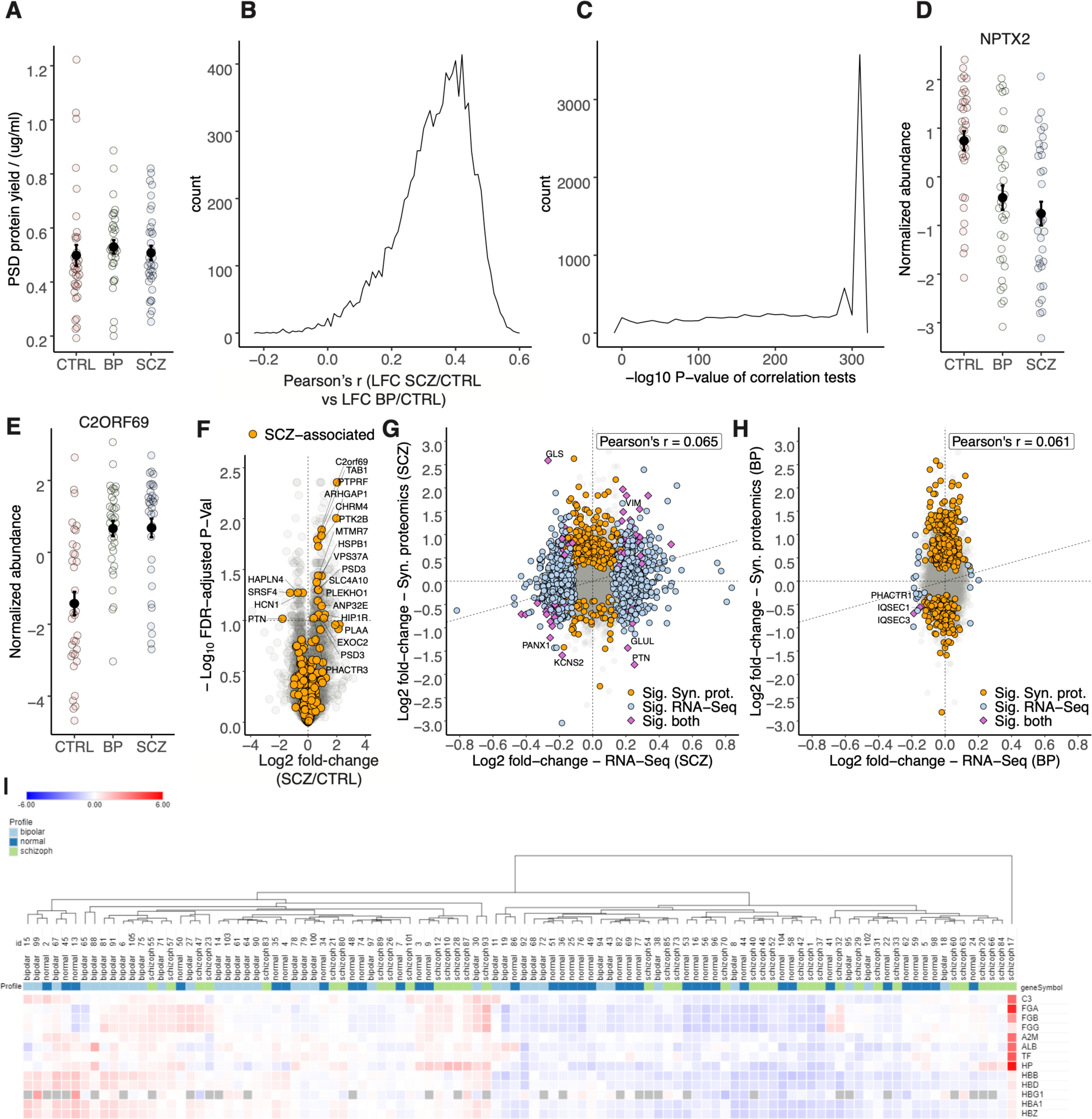
SCZ and BP synapses show substantial and similar proteomic changes. **(A)** Synaptic protein yields stratified by disease status. **(B)** CTRL samples were split in half, and each half was used to compute SCZ/CTRL and BP/CTRL LFCs. Histograms showing Pearson’s correlations coefficients between the LFCs in SCZ and those in BP and **(C)** P-Values (-log10 transformed) of positive correlation tests observed in 10,000 trials of this exercise. Correlation test p-values equaling zero were replaced with 5E-324, which is reflected in the peak at the right tail of the histogram. Normalized abundance observed in the MS/MS data in **(D)** NPTX2 and **(E)** C2orf69 proteins, stratified by disease status. **(F)** Volcano plot of alterations in SCZ synapses; proteins encoded by genes linked to SCZ by GWAS are highlighted, and significantly altered proteins are labeled. Evaluation of alterations in RNA-Seq from the ACC against those observed in synaptic proteomics of the DLPFC in the same cohort of individuals in **(G)** SCZ, and **(H)** BP. **(I)** Heatmap showing the synaptic proteome levels of several blood-enriched marker proteins across individual CTRL, SCZ, and BP samples; columns are hierarchically clustered. The excluded sample (schizoph17) segregates out.

**Figure S3:**
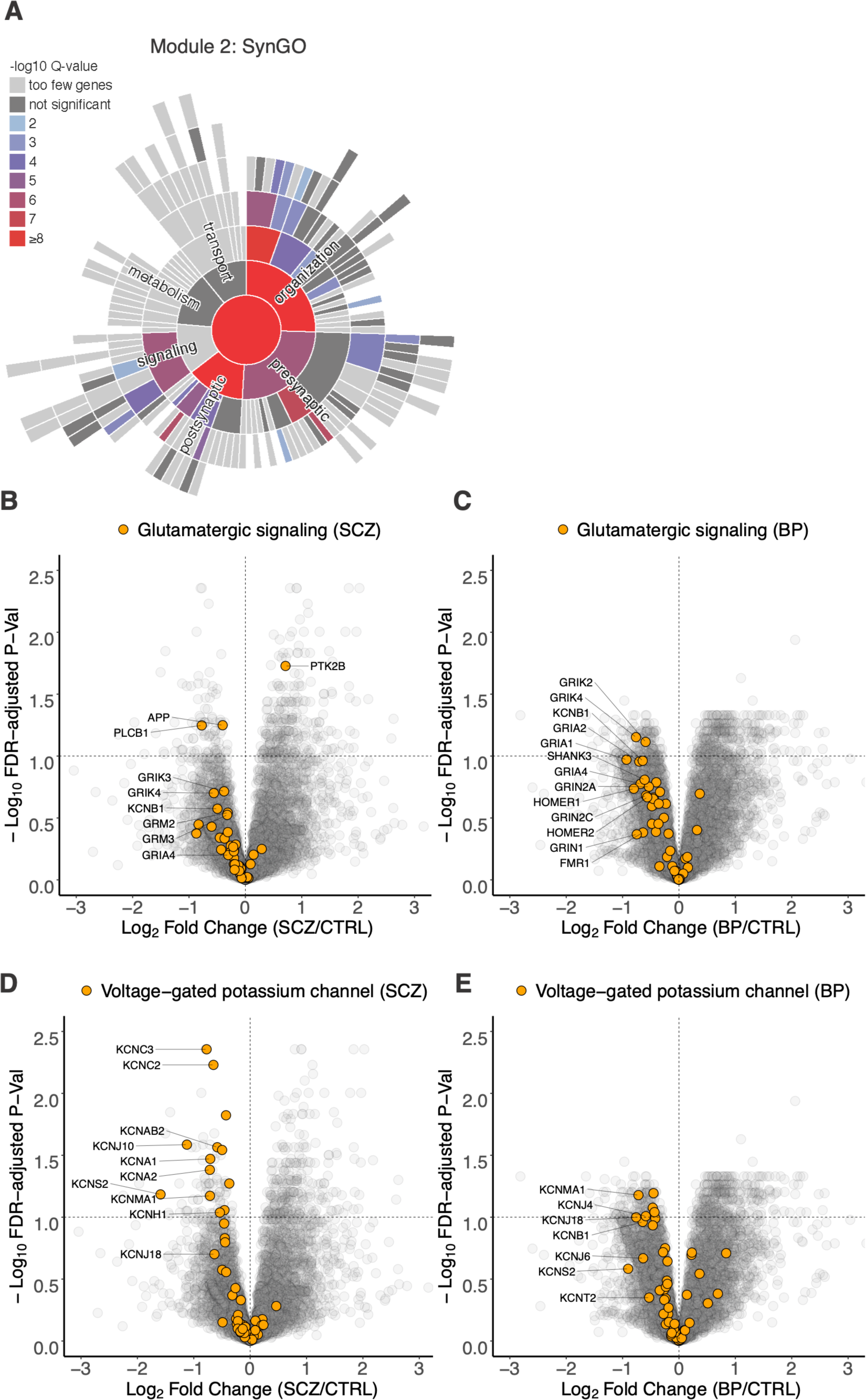
Core synaptic proteins are reduced in synapses from humans with SCZ and BP. **(A)** SynGo sunburst plot showing enrichment of biological processes related to synaptic organization, function, and signaling in module 2. Volcano plot showing alterations in **(B)** SCZ and **(C)** BP synapses, with all detected proteins in the ‘glutamate receptor signaling pathway’ in mSigDb highlighted. Volcano plot showing alterations in **(D)** SCZ and **(E)** BP synapses, with all detected proteins in the ‘voltage-gated potassium channel’ gene set in mSigDb highlighted.

**Figure S4:**
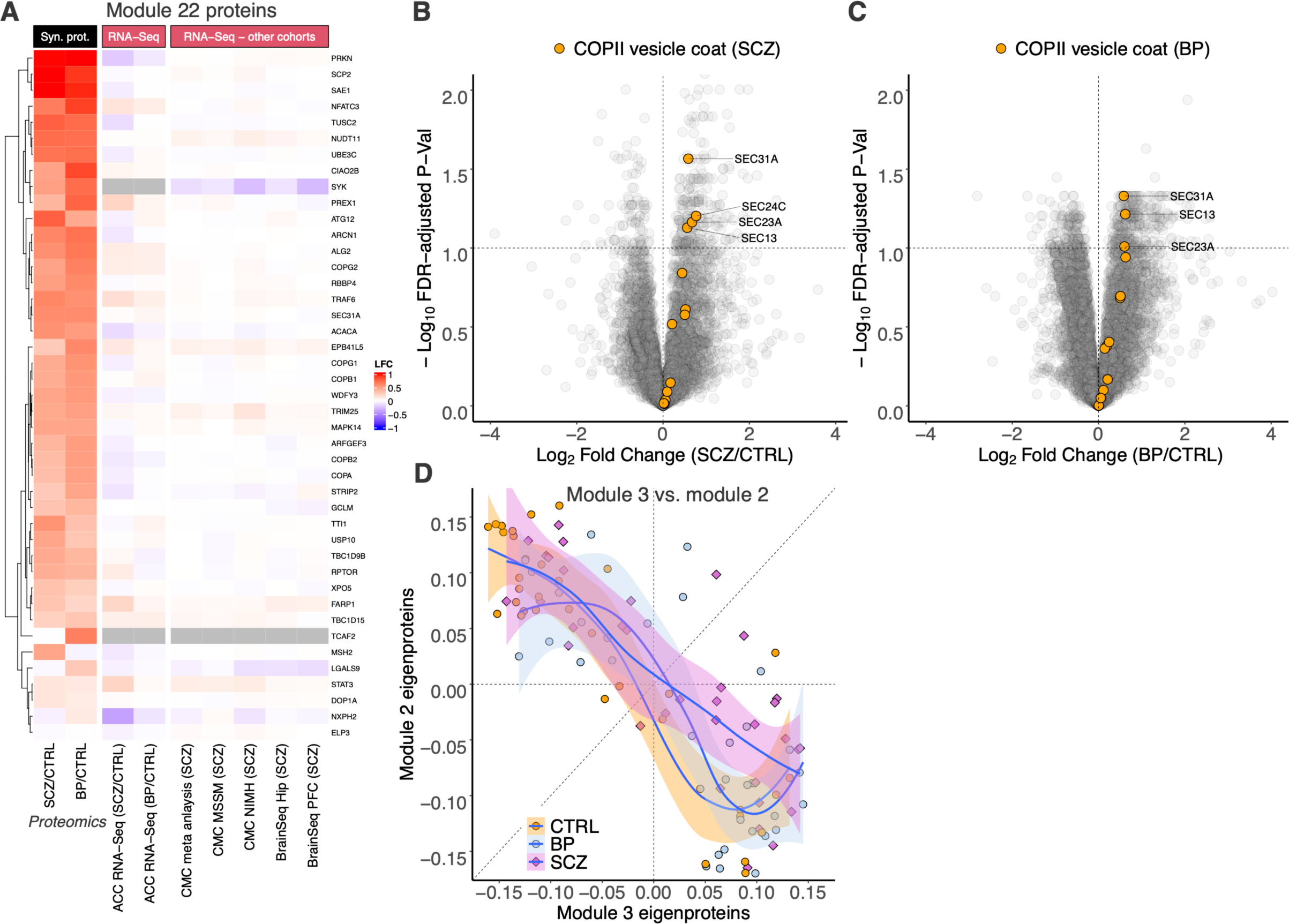
Proteins related to vesicle tethering and trafficking are upregulated in SCZ and BP synapses. **(A)** Heatmap depicting LFCs in all proteins in module 22. Heatmaps are hierarchically clustered by LFC in the synaptic proteome in SCZ and BP. RNA expression alterations (LFCs) in SCZ/BP obtained from previously described or publicly available datasets are shown alongside. Volcano plot highlighting alteration of COPII coat proteins in **(B)** SCZ, and **(C)** BP synapses. **(D)** Evaluation of WGCNA module 2 eigenproteins against those in module 3 in human post-mortem synaptic protein network. Eigenproteins are stratified by disease status and a local polynomial regression line fit to highlight correlation.

**Figure S5:**
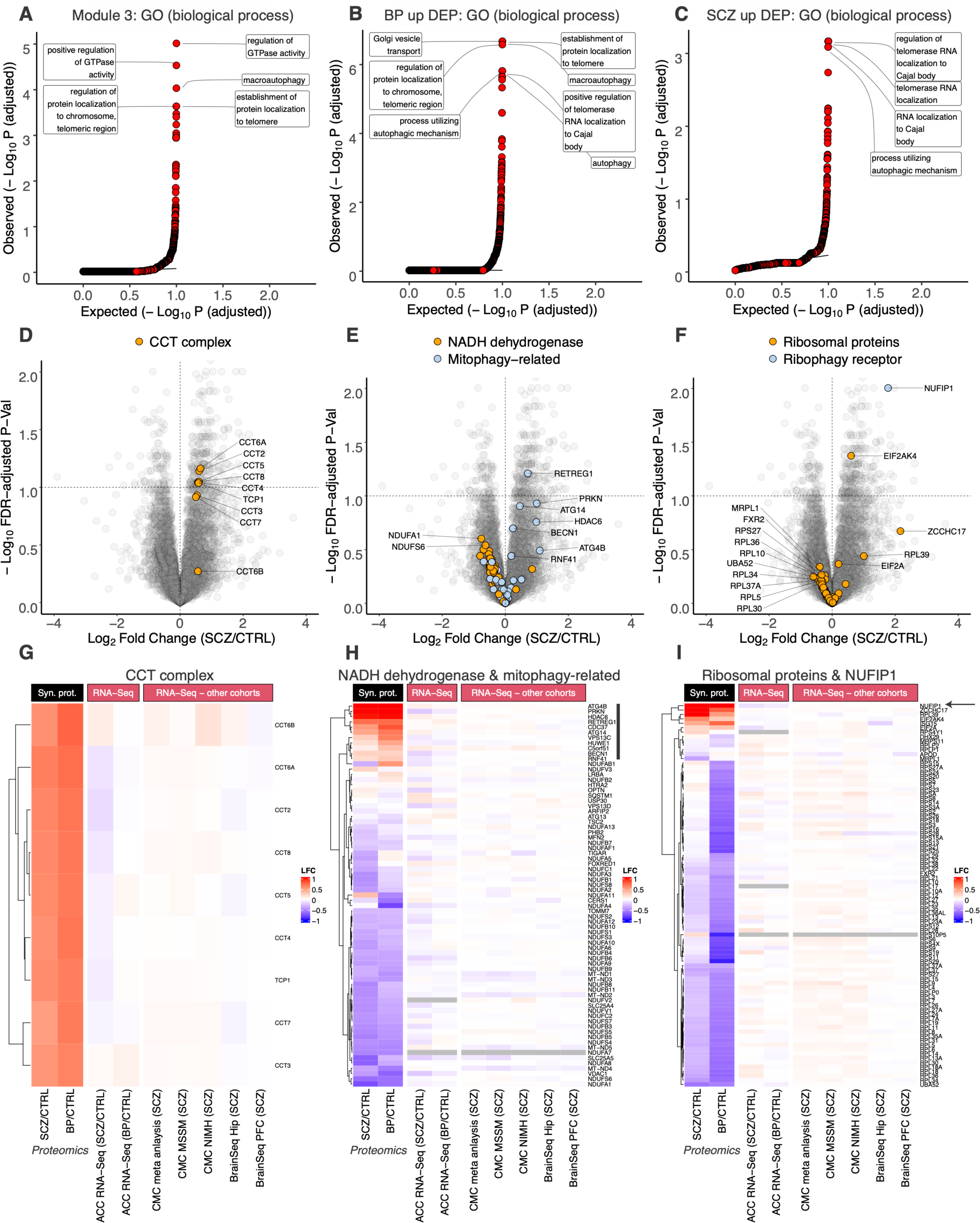
Autophagy pathways are elevated in SCZ and BP synapses. Q-Q plot showing biological processes enriched in **(A)** module 3, **(B)** upregulated DEPs in BP synapses, and **(C)** upregulated DEPs in SCZ synapses. The enrichments for telomerase-related gene sets are primarily driven the CCT complex proteins. Volcano plot showing alterations in SCZ synapses highlighting all detected proteins in the **(D)** ‘chaperonin-containing T complex,’ **(E)** ‘NADH dehydrogenase complex,’ and ‘mitophagy’, and **(F)** ‘cytosolic ribosome’ gene sets in mSigDb. Also highlighted is the ribosome receptor for ribophagy NUFIP1. Heatmaps depicting LFCs in all detected proteins in the **(G)** ‘chaperonin-containing T complex,’ **(H)** ‘NADH dehydrogenase’ and ‘mitophagy’ **(I)** ‘cytosolic ribosome’ gene sets in mSigDb. The ribophagy receptor NUFIP1 is shown as well and labeled with an arrow. Heatmaps are hierarchically clustered by LFC in the synaptic proteome in SCZ and BP. RNA expression alterations (LFCs) in SCZ/BP obtained from previously described or publicly available datasets are shown alongside. Thick black line in (H) labels the proteins in the mitophagy gene set upregulated in SCZ and BP synapses.

**Figure S6:**
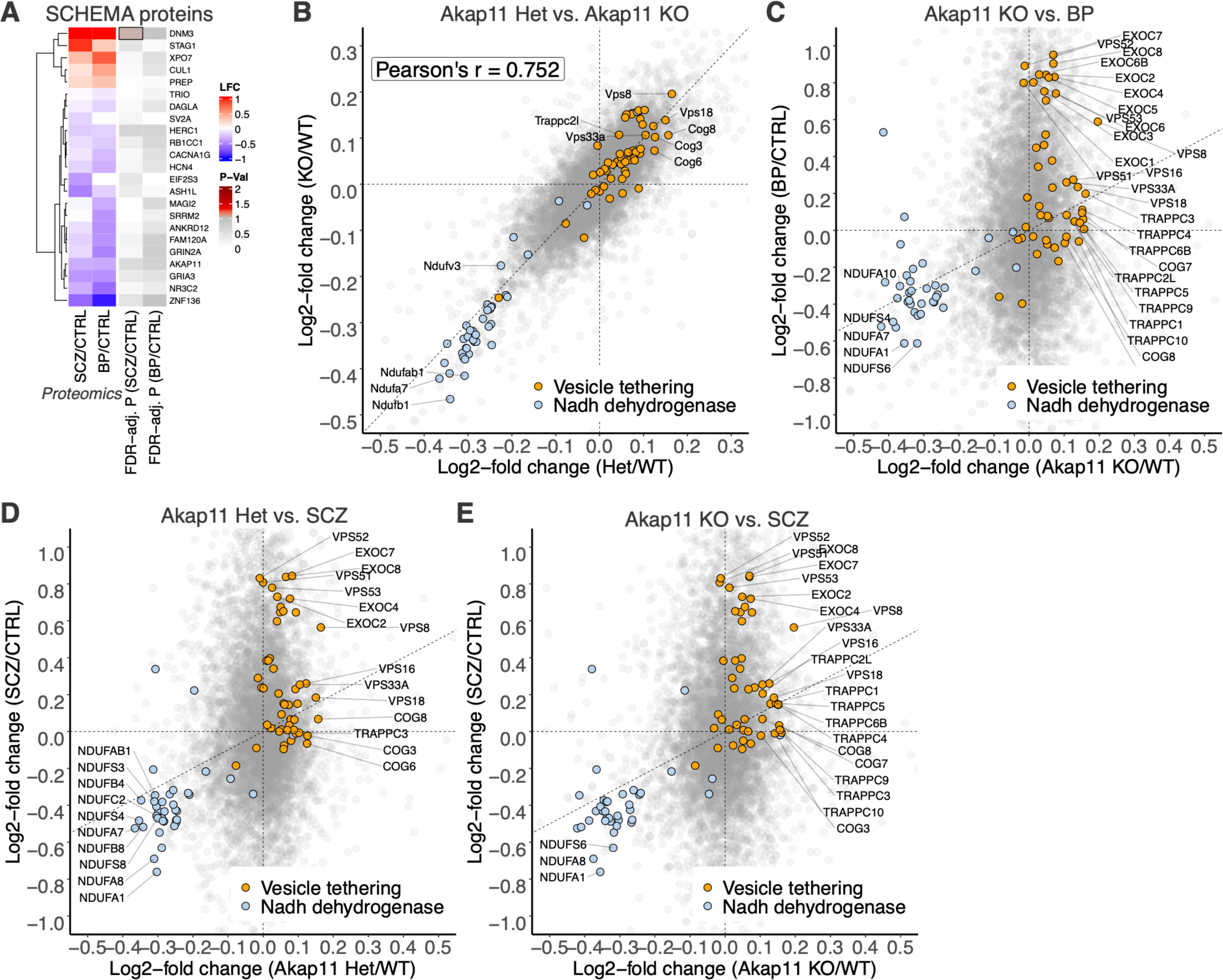
Comparison of alterations in SCZ and BP synapses with those in *Akap11* mice. **(A)** Heatmap showing LFCs in detected proteins of genes identified by the SCHEMA study as SCZ risk factors. Synaptic proteomics FDR-adj. P-values for SCZ/CTRL and BP/CTRL comparisons (-log10 transformed) are shown alongside; cells for DEPs (FDR-adj. P < 0.1) are outlined. **(B)** Comparison of the alterations (LFCs) in *Akap11* Het synapses against those in KO; gene sets related to mitochondrial respiration (‘NADH dehydrogenase complex’) and vesicle trafficking (‘vesicle tethering complex’) are colored. Loss of one of *Akap11* substantially recapitulates the effects observed with loss of both copies. **(C)** Evaluation of the LFCs in *Akap11* KO synapses against those in BP synapses; gene sets related to mitochondrial respiration (‘NADH dehydrogenase complex’) and vesicle trafficking (‘vesicle tethering complex’) are colored. **(D)** Same as (C) but showing *Akap11* Het vs. SCZ. **(E)** Same as (B) but depicting *Akap11* KO vs. SCZ.

### Methods

#### Human brain tissue samples

Human post-mortem dorsolateral prefrontal cortex (DLPFC; Brodmann’s Area 46) was obtained from the Stanley Medical Research Institute (SMRI). The analyzed ‘Array Collection’ consists of 105 subjects (35 control cases, 35 schizophrenia and 35 bipolar disorder patients; further demographic information available in Supplementary Table T1). One BP sample was excluded from the analysis based on advice from the SMRI, whose pathologist observed evidence of brain atrophy in the sample. Furthermore, one SCZ sample was excluded based on evidence of blood contamination in the brain tissue (see Fig. S1I). Information on prescribed medication and cause of death can be acquired from the SMRI. Investigators were blinded to group identity until completion of the proteomics data analysis.

#### Mouse samples

Whole cortices from one-month old male *Akap11* mutant mice and control littermates (n=5 wild-type (WT), 5 heterozygous knockouts (Het), 5 homozygous knockouts (KO)) were used in this study. Mutant mice were obtained from the Jackson Laboratory (B6.Cg-*Akap11^tm1.2Jsco^*/J ; Strain #: 028922). Exons 6 and 7 of the mouse *Akap11* gene have been deleted in this mutant to create a knockout allele. Mice were euthanized using CO_2_ inhalation, after which whole cortices were dissected and flash-frozen in liquid nitrogen.

#### Purification of synaptic fractions

Human and mouse synapses were isolated as previously described (Dejanovic et al., 2018). Briefly, flash-frozen DLPFC gray matter (human) or whole cortex (mice) was dounce-homogenized in ice-cold homogenization buffer (5 mM HEPES pH 7.4, 1 mM MgCl_2_, 0.5 mM CaCl_2_, supplemented with phosphatase and protease inhibitors). The homogenate was centrifuged for 10 minutes at 1,400 g (4°C) and the supernatant was re-centrifuged at 13,800 g for 10 minutes (4°C). The resulting pellet was resuspended in 0.32 M Sucrose, 6mM Tris-HCl (pH 7.5) and layered gently on a 0.85 M, 1 M, 1.2 M discontinuous sucrose gradient (in 6mM Tris-HCl pH 7.5) and ultracentrifuged at 82,500 g for 2 hours (4°C). The synaptosome fraction—which sediments at the 1 M and 1.2 M sucrose interface—was collected, an equal volume of ice-cold 1% Triton X-100 (in 6mM Tris-HCl pH 7.5) was added, mixed thoroughly, and incubated on ice for 15 minutes. The mixture was ultracentrifuged at 32,800 g for 20 minutes (4°C), and the final synapse protein pellet was collected by resuspension in 1% SDS. A small aliquot was taken to measure the protein concentration using the BCA assay (ThermoFisher Scientific), and the remaining protein was stored at -80°C until being processed for mass spectrometry.

#### Processing of human samples for Mass Spectrometry (MS)

Synaptic fractions were purified from the DLPFC of 105 CTRL, SCZ, and BP post-mortem human brains. The group designations were blinded as A, B and C with 35 samples in each category. Samples containing 30 to 40 µg protein pre-digestion were processed in batches of 15 (with 5 samples from each group) using the following proteomic workflow.

All samples were prepared in 1% SDS to ensure proper dissolution of membrane proteins and digested using S-Trap^TM^ sample processing technology (Protifi) following manufacturer’s instructions. Reduction of proteins using 5 mM dithiothreitol and alkylation using 10 mM iodoacetamide was performed at room temperature. The denatured, non-digested proteins were then processed on a micro s-trap column. The proteins were bound to the column via centrifugation and contaminants/detergents were washed away. Sequential digestion steps were then performed on column using 1:20 enzyme to substrate ratio of Lys-C for 2 hours and Trypsin overnight at room temperature.

Post-digest nanodrop measurements were performed and 14 µg aliquots per sample were made for labeling. A pooled reference sample was also created using equal amounts of all 105 samples. Seven tandem mass tag 16 (TMT16) plexes were constructed by first randomly assigning 5 samples from each group (A, B, and C) to a plex and then randomly designating channels to samples within a plex. The last channel 134N contained the pooled reference sample in all plexes. Samples were reconstituted in 50 mM HEPES buffer for labeling and 20 µL of 25 µg/µL TMT16 reagent was added for the labeling reaction. After confirming successful labeling (>95% label incorporation), the reactions were quenched with 5% hydroxylamine and combined. The mixed sample was then desalted on a 50 mg tC18 SepPak column and 224 µg of TMT16 labeled peptides were fractionated by high pH reversed-phase chromatography on a 2.1 mm x 250mm Zorbax 300 extend-c18 column (Agilent). One-minute fractions were collected during the entire elution and fractions were concatenated into 12 proteome fractions for LC-MS/MS analysis.

#### Processing of mouse samples for MS

Protein from purified synaptic fractions was digested as described above. 100 µg of each sample was labeled with a TMT16 reagent following the protocol described above. The combined sample was fractionated on a 4.6 mm x 250 mm Zorbax 300 extend-c18 column (Agilent) and concatenated into 18 fractions for LC-MS/MS analysis.

#### Liquid chromatography (LC) - MS/MS

One microgram of each fraction of human and *Akap11* mouse study was analyzed on a QE HFX mass spectrometer (ThermoFisher Scientific) coupled to an easy-nLC 1200 LC system (ThermoFisher Scientific). Samples were separated using 0.1% Formic acid / 3% Acetonitrile as buffer A and 0.1% Formic acid /90% Acetonitrile as buffer B on a 27 cm 75 µm ID picofrit column packed in-house with Reprosil C18-AQ 1.9 µm beads (Dr Maisch GmbH) with a 90 min gradient consisting of 6-20% B in 62 min, 20-30% B for 22 min, 30-60% B in 9 min, 60-90% B for 1 min followed by a hold at 90% B for 5min. The MS method consisted of a full MS scan at 60,000 resolution and an AGC target of 3e6 from 350-1800 m/z followed by MS2 scans collected at 45,000 resolution with an AGC target of 5e4 with a maximum injection time of 105 ms and a dynamic exclusion of 15 seconds. The isolation window used for MS2 acquisition was 0.7 m/z and 20 most abundant precursor ions were fragmented with a normalized collision energy (NCE) of 29 optimized for TMT16 data collection.

#### Database search and MS/MS quantification

The data was searched on Spectrum Mill MS Proteomics Software (Broad Institute) using either a human or mouse database that contained 69062 and 47069 entries downloaded from the UniProt database (The UniProt Consortium, 2021) on 08/05/2020 and 12/28/2017 respectively. The Spectrum Mill generated proteome level export which was filtered for proteins identified by two or more peptides for further analysis. Protein quantification was achieved by taking the ratio of TMT reporter ions for each sample over the TMT reporter ion for the pooled reference channel for the human study or the median of all channels for the *Akap11* mouse study. TMT16 reporter ion intensities were corrected for isotopic impurities in the Spectrum Mill protein/peptide summary module using the afRICA correction method which implements determinant calculations according to Cramer’s Rule (Shadforth et al., 2005) and correction factors obtained from the reagent manufacturer’s certificate of analysis (https://www.thermofisher.com/order/catalog/product/90406) for lot numbers VE299607 and VH310017 for human and mouse datasets, respectively.

#### Human dataset statistical analysis - linear modeling and regression of unwanted covariates

The database search and quantification process produces a dataset containing abundance measurements for proteins observed in each sample in the form of log-transformed ratios to the common reference present in each TMT plex. This data was normalized on a sample-by-sample basis using median centering followed by median absolute deviation (MAD) scaling. The normalized data was filtered to eliminate proteins that were missing values in more than 80% of the samples.

Examination of covariates using the metadata provided by the brain bank showed that multiple covariates had varying distributions among the three groups. These disparities between the groups, especially those in ‘cause of death,’ were also observed to contribute substantially to the variance within the dataset. To adjust for this covariate, a new ‘rate of death’ variable was created by binning cause of death into four bins as follows.

| Rate of death | Cause of death |
| --- | --- |
| 1 | Sudden (e.g., cardiac) |
| 2 | Possible anoxia at death (e.g., CO poisoning or pneumonia) |
| 3 | Slower death (e.g., death from cancer) |
| 4 | On mechanical ventilator prior to death |

Differentially expressed proteins for the SCZ/CTRL and BP/CTRL comparisons were determined using a linear model which adjusted for rate of death, sex, and post-mortem interval. The linear model was implemented using the *limma* library in R (Ritchie et al., 2015). Nominal p-values obtained using the linear model for each protein were adjusted for multiple testing using the Benjamini-Hochberg FDR approach (Benjamini and Hochberg, 1995).

#### Statistical analysis of the Akap11 mouse dataset

After performing median-MAD normalization, a moderated two-sample t-test was applied the datasets to compare WT, Het and KO sample groups. Benjamini-Hochberg corrected p-value thresholds were used to assess differentially expressed proteins between experimental conditions.

#### Overlap with GWAS hits

GWAS risk genes for neurological and psychiatric disorders and behavioral traits were obtained from Yang et al. (2021). Gene symbols for each disorder/trait were processed to include only protein coding genes with annotations in the UniProt database (The UniProt Consortium, 2021). Specifically, gene symbols starting with “LOC”, “LINC”, “ENSG”, “MIR” were removed, as were gene symbols composed entirely of numbers, those with a period or a space within the symbol, and those with the symbol “NA”. Duplicated gene symbols were cleaned up so that only one instance of the symbol was present. In the rare occasions where two gene symbols were present in the same cell separated by a “/”, both gene symbols were considered.

#### RNA-Seq data analysis

Paired-end RNA-Seq data were obtained from the SMRI in *fastq* format. RNA-Seq methodology is described in (Hwang et al., 2013) and (Ramaker et al., 2017). Data for a total of 82 samples (n = 26 CTRL, 25 BP, 31 SCZ) was provided to us. Sequencing adapters were trimmed using *Trimmomatic* (Bolger et al., 2014). The surviving reads were aligned to GRCh38 primary assembly using *STAR* (Dobin et al., 2013). Reads aligning to transcript regions annotated in the GENCODE ‘comprehensive’ gene annotation were counted using *featureCounts* in the ‘unstranded’ mode to obtain gene-level counts (Liao et al., 2014). The counts for all samples were collated into a table and used as input for *DESeq2* for statistical analysis (Love et al., 2014). No outliers were detected after principal component analysis of the normalized, variant-stabilized count matrix. To control for confounders, covariates including age, sex, post-mortem interval, and brain pH were included in the design formula, alongside the primary variable of interest (disease status). Only genes with at least 5 counts in 75% of the samples were included in the analysis. Dispersions and log2 fold changes were estimated using the function *DESeq*, and differential expression was inferred using Wald test p-values. The threshold for differential expression was set at a false discovery rate (FDR) of 0.1.

#### Gene set enrichment analysis (GSEA)

GSEA was performed with the R Bioconductor package *fgsea* (Sergushichev, 2016), using C5 ontology gene sets obtained from the Molecular signatures database (MSigDB) (Liberzon et al., 2011). Proteins were pre-ranked using their linear model test-statistics within each comparison (SCZ/CTRL or BP/CTRL), and their gene symbols were used for the analysis. For proteins with multiple isoforms (n=551 of 8,996), the isoform with the largest effect-size was used for pre-ranking. Because the enrichment analyses were carried out using the proteins’ gene symbols, we use the terms gene set and protein set interchangeably in this study. Gene sets with FDR-adjusted p-values < 0.1 were considered significant.

#### Network analysis

Network analysis was performed using WGCNA (Langfelder and Horvath, 2008). The raw data matrix was composed of log2-transformed TMT ratios that were zero-centered and scaled using the median-MAD (median absolute deviation) approach. 887 proteins were removed because they had too many missing values or zero variance; no outlier samples were detected. A weighted protein co-expression network was generated using the sample (n=103) X log2 ratio (n=8079) matrix. A soft-threshold power of 5 was chosen because (1) the signed scale-free topology model fit (R2) approached an asymptote of ∼0.8 at that threshold, and (2) the mean connectivity at that power was just under 100. Satisfying both these requirements approximates scale-free topology in the network.

We used the *blockwiseModules* function to generate the network, with the settings soft- threshold power = 5, minimum module size = 25, network type = “signed”, deep split = 4, and merge cut height = 0.15. The correlation type used was Pearson’s r, and the TOM denominator was set to ‘mean.’ The partitioning around medoids-like (PAM-like) stage of module detection was enabled, with proteins being PAM-assigned only to clusters that lie below it on the branch that it was merged into (*pamRespectsDendro* = TRUE). The maximum block size was set at 10,000 to complete all clustering in a single block. Briefly, using this approach, we calculated pairwise Pearson’s correlations between all possible protein pairs and obtained a 8079 X 8079 correlation matrix. This co-expression similarity (correlation) matrix was transformed into a signed adjacency matrix by raising it to the soft-threshold power of 5. The signed adjacency matrix was used to generate a topological overlap matrix (TOM), which represents pattern similarity across protein pairs, and serves as an important filter against spurious or missing connections between network nodes (Yip and Horvath, 2007). Hierarchical clustering was carried out on TOM-based dissimilarity (1-TOM), and the resultant dendrogram was used for module identification. Modules were identified using the dynamic branch cutting algorithm, as well as a second stage of PAM- like partitioning, implemented within the *blockwiseModules* function.

#### Merging datasets

*Homo sapiens* homolog-associated gene symbols were obtained for all mouse genes from Ensembl BioMart. Rows from the human and *Akap11* synaptic proteomics datasets were merged conditioned on matching homologous gene symbols. For proteins with more than one isoform, the isoform with the largest number of spectra was chosen. The same rule was also used to merge synaptic proteome and RNA-Seq datasets.

#### Gene ontology (GO) analysis

GO analysis was carried out using the R library *clusterProfiler*. Human cellular component, biological process, and molecular function GO terms were obtained from the Bioconductor OrgDb org.Hs.eg.db. Over-representation analysis was carried out using the function *enrichGO.* The subset of uniquely detected proteins that were used to construct the WGCNA network (n=8079) were used as the background. Adjusted p-values (q-vals) reported by *enrichGO* were used to make Q-Q plots to discern gene sets deviating from the uniform distribution.

#### Assigning biological pathway to module

GO analysis of module proteins was performed using org.Hs.eg.db gene ontologies and GSEA performed using mSigDb C5 gene sets. Unique gene symbols of proteins in each module were used for either analysis. Convergent signal originating from GO terms significant in cellular component GO analysis (FDR-adj. P < 0.1) and GSEA (gene sets FDR-adj. P < 0.1) were prioritized, after which significant biological process and molecular function GO terms were used to define modules’ principal biology.

#### Data availability

The original mass spectra and the protein sequence databases used for searches have been deposited in the public proteomics repository MassIVE (http://massive.ucsd.edu) and are accessible at ftp://MSV000090320@massive.ucsd.edu with the password: schizophrenia. If requested, also provide the username: MSV000090320. The datasets will be made public upon acceptance of the manuscript.

